# Age-dependent Dysferlin Accumulation in Macrophages Promotes STAT1 Activation via Calcium Influx, Impairing Myogenesis

**DOI:** 10.1101/2025.10.02.679938

**Authors:** Kana Tominaga, Naoomi Tominaga

## Abstract

DYSF functions as a regulator of Ca²⁺ in the skeletal muscle and facilitates muscle repair following injury. It is also highly expressed in monocytes and macrophages and related to inflammation, immune regulation, and the mononuclear phagocyte system. Macrophages are pivotal in driving myogenesis, and specific cytokine-induced macrophage differentiation plays a role in maintaining skeletal muscle during disease and aging. Thus, a comprehensive understanding of the mechanisms by which DYSF operates in macrophages may inform the development of novel treatments for muscle atrophy. In this study, we demonstrated that DYSF expression is associated with monocyte differentiation and increases with age. Our findings indicate that DYSF overexpression induces the generation of M1-type macrophages, which subsequently secrete inflammatory cytokines and promote cell invasion. Furthermore, we observed that DYSF regulates Ca²⁺ influx into the cell and activates the STAT1 signaling pathway, whereas DYSF deficiency suppresses these processes. Macrophages overexpressing DYSF are associated with the inhibition of myoblast differentiation during myogenesis in a co-culture system involving macrophages and myoblasts. Therefore, the balance between the STAT1 signaling pathway and Ca²⁺, which are regulated by DYSF abundance, may play a crucial role in myogenesis.

## Introduction

Dysferlin (DYSF) is a type II transmembrane protein located in the plasma membrane. It comprises two DysF domains and seven C2 domains (C2A-C2G, with 13%-33% identity), which function as calcium ion (Ca²⁺)-binding modules (*1*, *2*). DYSF is crucial for membrane repair, which is dependent on extracellular Ca²⁺ (*1*, *3*). Following injury, Ca²⁺ influx into the cell binds to the C2 domains of DYSF, facilitating the fusion of nearby vesicles to seal the injury site. DYSF is also localized at the T-tubules in the skeletal muscle and may regulate Ca²⁺ release from the sarcoplasmic reticulum (*4*). The absence of DYSF results in persistent Ca²⁺ influx, membrane injury, exacerbating fiber necrosis, activating proteases such as calpains, and leading to mitochondrial dysfunction and cell death (*5*, *6*). In patients with dysferlinopathy caused by pathogenic mutations in the DYSF gene, DYSF protein is not localized to the cell membrane, resulting in impaired membrane repair when the skeletal muscle is damaged (*7*).

DYSF is also highly expressed in monocytes and macrophages at levels comparable to those in skeletal muscles (*8*). In patients with dysferlinopathy, peripheral blood monocytes serve as a less invasive diagnostic method because of the correlation between expression levels and genetic mutations in monocytes and skeletal muscle (*9*, *10*). DYSF expression is upregulated in differentiating monocytes in the THP1 monocyte cell model (*11*). In monocytes, DYSF regulates vesicle trafficking and phagocytosis, which are Ca²⁺-dependent processes (*12*, *13*). DYSF-deficient monocytes produce more inflammatory signals than normal monocytes (*14*). It has been indicated that DYSF cooperates with integrin beta3 (ITGB3) and focal adhesion components to enhance monocyte adhesion capacity (*11*). Loss of DYSF induces ITGB3 expression, thereby increasing cell motility. These processes may contribute to early inflammation in the muscle before extensive degeneration occurs. *Dysf*-deficient muscles exhibit massive macrophage infiltration and expansion, even without overt injury. However, in *Dysf*-deficient BLA/J mice, macrophages accumulate in the muscle regardless of DYSF expression (*15*). *Dysf*-deficient mice transplanted with healthy DYSF-expressing macrophages showed only mild benefits in muscle regeneration (*16*). These reports indicate that the function of DYSF-expressing macrophages involved in myogenesis has not yet been fully elucidated.

Maintaining muscle strength necessitates skeletal muscle regeneration, which encompasses not only the repair of the sarcolemma by membrane repair-related proteins such as DYSF but also the formation of muscle fibers through the proliferation, differentiation, and fusion of skeletal muscle stem cells. In the context of skeletal muscle regeneration, interactions with various cells surrounding muscle cells are critical. Macrophages are implicated in diseases and aging and are characterized by a propensity for excessive inflammatory responses in various tissues. Monocytes migrate to tissues and differentiate into macrophages, which primarily exist in two states: M1- and M2-type (*17*). M1-type macrophages are identified by the secretion of inflammatory cytokines, such as tumor necrosis factor–α (TNF-α) and Interleukin-6 (IL-6), and the activation of inducible nitric oxide synthase (iNOS) and the signal transducer and activator of transcription 1 (STAT1) by IFN-gamma (IFN-γ) or lipopolysaccharide (LPS), inducing excessive inflammatory responses in various tissues (*18*). M2-type macrophages are characterized by the expression of IL-10, mannose receptor (Mrc1), and Arginase-1 (Arg-1) activated by IL-4, and are involved in the fusion stage of skeletal muscle fibers. In aging skeletal muscle, macrophage polarization is altered. Human studies report an increase in M2-type macrophages, whereas M1-type macrophages are reduced (*19*). M2-type macrophages often are co-localized in intermuscular adipose tissue (IMAT) (*20*). This shift may be driven by fibro-adipogenic progenitors (FAPs), which recruit and polarize macrophages via C-C motif chemokine ligand 2 (CCL2) and other secreted factors (*21*). In contrast, mouse studies show an accumulation of inflammatory macrophages with aging, accompanied by inflammation, oxidative stress, and activation of pro-inflammatory and extracellular matrix pathways (*22*). Despite species differences, aged muscle exhibits a pro-inflammatory status. These mechanisms may include “non-canonical” M2-type macrophage activation (producing TNF-α, IL-1β, IL-6) and contributions from other tissues such as adipose tissue, which release pro-inflammatory cytokines into circulation (*23*). Overall, in elderly individuals, the persistence of inflammatory M1-type macrophages after muscle injury may contribute to age-related muscle atrophy, while M2-type macrophages may contribute to fibrosis and IMAT development. Both M1- and M2-type macrophages constitute the major infiltrating macrophage phenotypes in the skeletal muscle of DYSF-deficient Bla/J mice (*24*). The presence of both macrophage types indicates an active inflammatory environment in the muscle attempting to repair damage; however, it remains unclear whether DYSF in macrophages is crucial for this process.

We here investigated how DYSF expression in macrophages changes with age. The research proponent compared DYSF expression levels and function in macrophages from young to elderly individuals and mice. We found that DYSF expression was higher in the elderly than in the young, indicating the characterization of macrophage polarization. DYSF overexpression promotes the activation of IFN-γ/STAT1 signaling via an increase in Ca²⁺ concentration and the secretion of inflammatory cytokines, such as TNF-α and IL-6. Furthermore, we found that M1-type macrophages induced by Ca²⁺ signaling suppressed the differentiation and fusion of skeletal muscle cells. Given that Ca²⁺ signaling is associated with aging, it represents a potential therapeutic strategy for age-related atrophy with high DYSF expression levels.

## Results

### DYSF is high-expression Monocytes during Aging

The role of DYSF in the differentiation of human monocytes into macrophages during aging was also investigated. Initially, we analyzed the gene expression data (GSE56047) (*25*) from human CD14-positive monocytes to explore the relationship between DYSF expression levels and aging (Fig. 1A). Our analysis revealed that DYSF mRNA expression levels increased in an age-dependent manner (*p*-value = 2.985e-09, R = 0.388) (Fig. 1B). To validate these findings, we isolated CD11b-positive monocytes from the bone marrow of young (8-week-old, 5 females/5 males) and aged (30-month-old, 5 females/5 males) mice. We observed that DYSF mRNA expression in CD11b-positive monocytes was upregulated in aged mice compared to that in young mice (Fig. 1C). Furthermore, there was a significant accumulation of DYSF protein in aged mice (Fig. 1, D and E), suggesting that aging may be associated with changes in the monocyte phenotype involving the DYSF protein.

**Figure 1.**
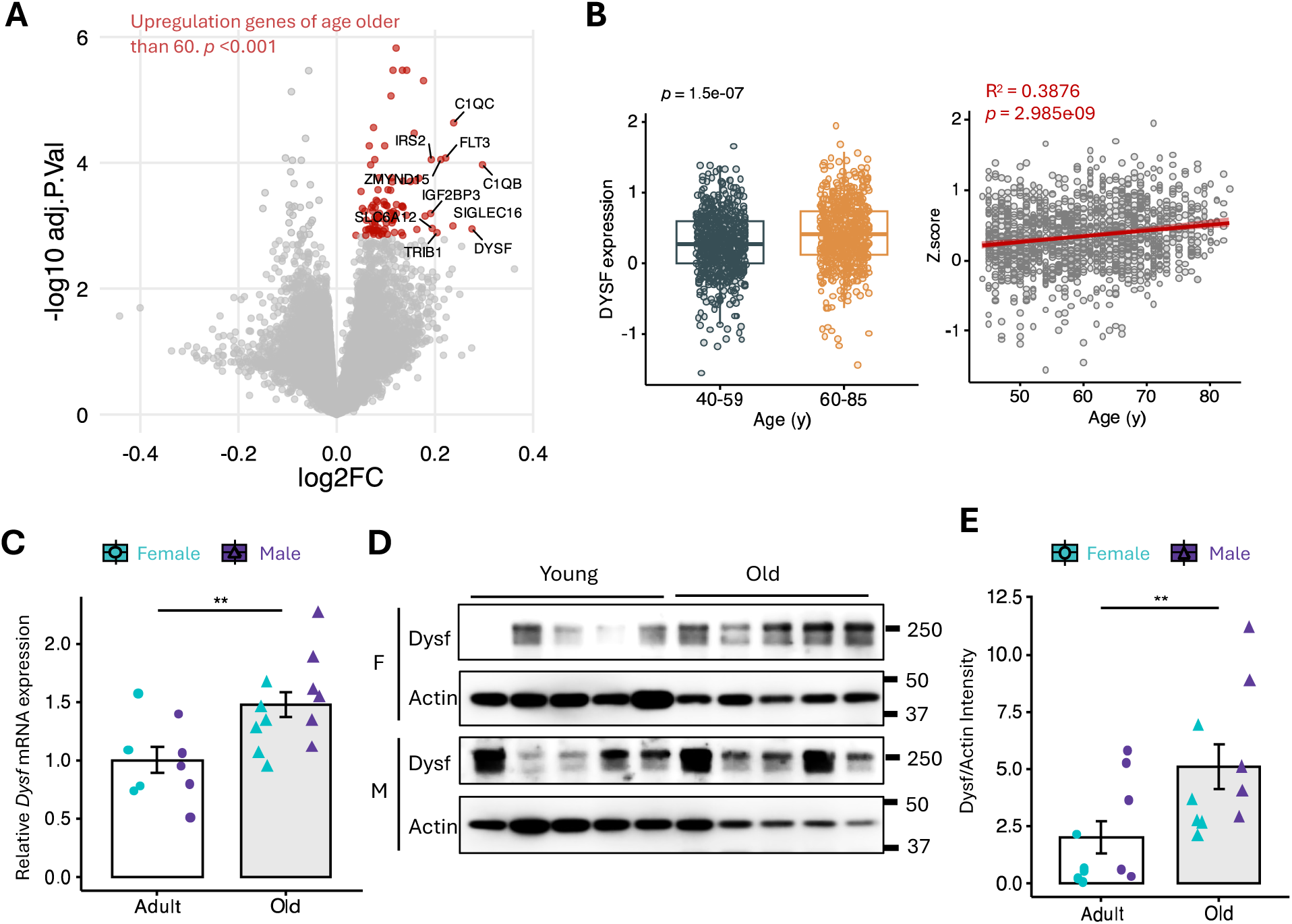
Aging-Related Increase of DYSF in Monocytes. (A) Volcano plot of the transcriptional microarray data (GSE56047) containing human peripheral CD14+ cells collected from 1,202 individuals ranging 44 - 83 years of age. (B) Scatter plot of DYSF gene expression between adult (44 – 59 years old, n = 593) and elder (60 – 83 years old, n = 609) subjects in GSE56047 database (left). Regression analysis between DYSF gene expression (Z-scores) and age (right). (C) Expression of DYSF mRNA was assessed by qPCR to compared adult (6 - 8 months) and old mice (30 months). Samples sizes of per group: adult female, n = 5; old female, n = 5; adult male, n = 5; old male, n = 5. (D) The protein levels of Dysf in 6-month-old (Adult) or 30-month-old (Old) mice were evaluated by western blotting. β-Actin (Actin) was used as the loading control. F, female; M, male. (E) Whole protein abundance of Dysf in 6-month-old (Adult) or 30-month-old (Old) mice. Band intensity was normalized to Actin. (B) (left), (C), (E), (G), and (H) were analyzed by unpaired Student’s t-test. (G) and (H) were analyzed by One-way ANOVA and Tukey-Kramer test. Figures [(C), (E), (G), and (H)] represented as means ± SD. *P < 0.05, ** P < 0.01, and *** P < 0.001.

Macrophage infiltration is a critical immune response whichein circulating monocytes migrate to tissues and differentiate into macrophages to perform essential repair functions and facilitate skeletal muscle regeneration. To examine whether DYSF is expressed during differentiation, we conducted a differential gene expression analysis to identify changes in the differentiation of monocytes. The phorbol 12-myristate 13-acetate (PMA)-induced U937, a human monocyte cell line, is widely used as an *in vitro* model for studying human macrophage function. Volcano analysis of the public microarray data (GSE107566) (*26*) showed that DYSF expression was upregulated by PMA-induced differentiation (fig. S1A). We observed an increase in DYSF mRNA expression in PMA-treated U937 cells compared to that in untreated cells in a time-dependent manner (fig. S1B). These results were further validated by flow cytometry analysis and immunostaining in U937 cells treated with PMA, confirming the accumulation of DYSF protein after post-PMA treatment (fig. S1C, D). These findings indicate that DYSF is increased during monocyte differentiation.

### DYSF is Accumulated in Aged Macrophages within Skeletal Muscle Tissue

Macrophages exhibit considerable diversity, and their functions and polarization range from M1- to M2-type states, playing a crucial role in effective muscle repair. This study focused on infiltrating macrophages in steady-state skeletal muscle, examining their role in skeletal muscle inflammation in both aged and young mice. At first, histological analyses confirmed the low density of myofibers in the skeletal muscle of aged mice (fig. 2A). Subsequent analyses involved assessing M1- or M2-type macrophage phenotypes and evaluating DYSF expression through immunofluorescence. Immunostaining with the CD80 marker revealed an increased accumulation of M1-type macrophages in aged skeletal muscle (fig. S2, A and B). Conversely, there was little differences in CD206-positive M2-type macrophages (fig. S2, C and D), suggesting a sustained proinflammatory state due to the accumulation of M1-type macrophages during aging. Immunostaining demonstrated the co-localization of DYSF with CD11b, with a higher presence of DYSF-positive macrophages in aged mice than in the young cohort (Fig. 2, B and C). These findings indicate that chronic low-grade inflammation is a characteristic of aging skeletal muscle, significantly influenced by increased pro-inflammatory macrophage infiltration associated with DYSF expression.

**Figure 2.**
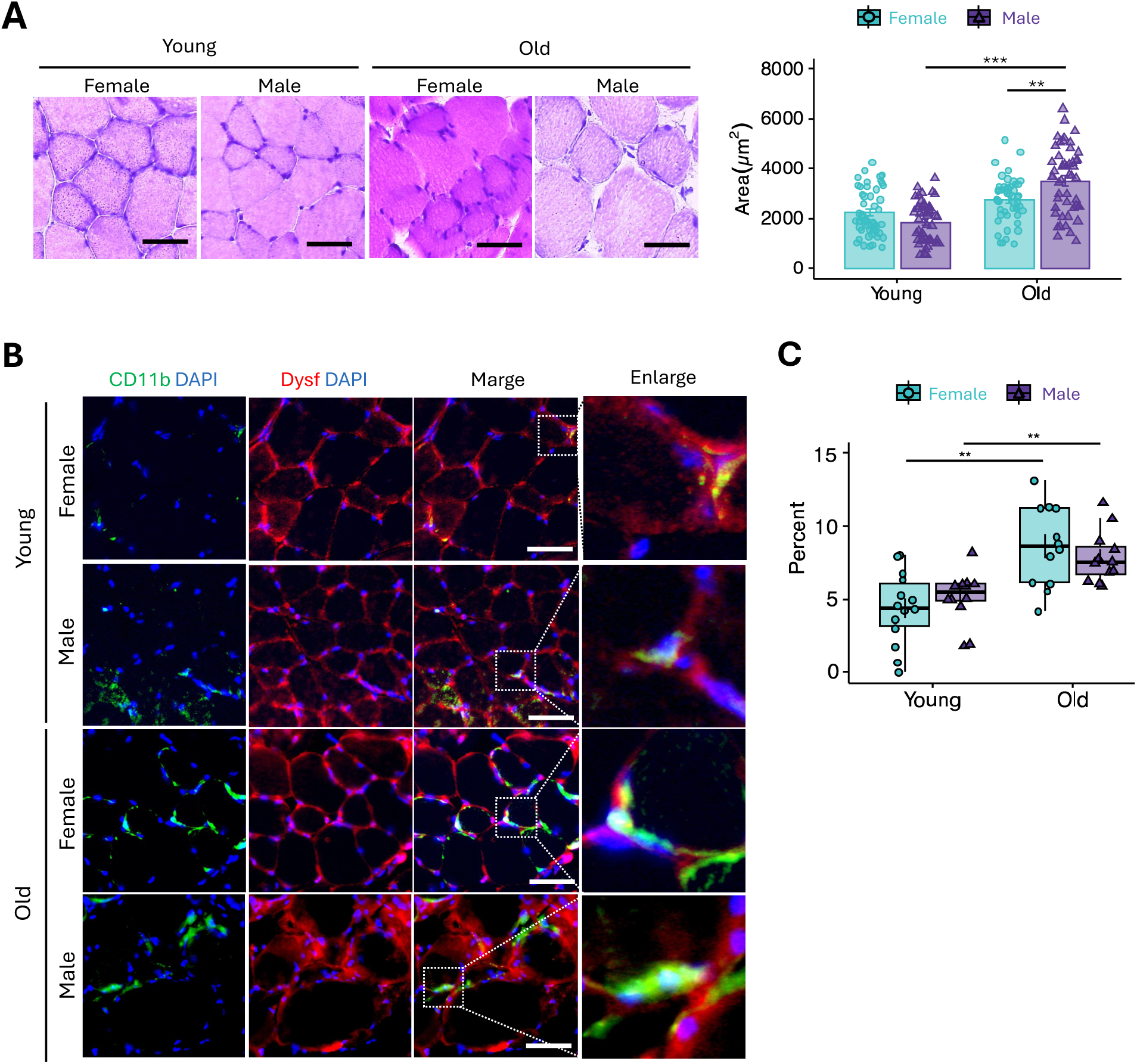
Dysf-expressing Macrophages increases in Age-related Skeletal Muscle. (A) Transverse sections of the soleus muscle were prepared and stained with Hematoxylin and eosin (HE) to assess muscle structure and morphology (left). Cross-sectional area (CSA) of muscle fibres (µm²) (right). n = 50. Scale bars, 25 μm. (B) Representative images of frozon sections of soleus muscle from 8-week-old (Young) and 30-month-old (Old) mice, stained with antibodies against CD11b and Dysf. DAPI was used as a nuclear stain. Scale bar, 50 µm. (H) Plot of image analysis data, showing the intensity per field measured for CD11b and Dysf double-positive cells. (A) (right) and (C) were analyzed by One-way ANOVA and Tukey-Kramer test, and represented as means ± SD. * P < 0.05, ** P < 0.01, and *** P < 0.001.

### DYSF Induces Polarization of M1-type Macrophages

We next examined whether high DYSF expression in macrophage was influence the polarization. We transfected constructs, DYSF plasmids (DYSF; DYSF-3HA) or control emptiy vector (Control; pcDNA3.1) into J774A.1, a murine monocyte/macrophage cell line (fig. S3, A and B). We then measured cell proliferation, cell area, aspect ratio and circularity to assess the charactarization of DYSF-overexpressing cells. DYSF-overexpressing cells showed a slight suppression of cell proliferation compared to control cells, exhibiting the characteristics of macrophage differentiation (fig. S2C). Cell area and aspect ratio increased with differentiation, while circularity decreases (*27*). We found that DYSF-overexpressing cells were larger in size, elongation and less circular compared with control (Fig. 2, A and B). We next performed knockdown experiments in J774A.1 using small interfering RNA (siRNA) for DYSF (fig. S4A). Cell area and aspect ratio were significantly decreased, and circularity was conversely increased by DYSF knockdown in J774A.1 under treatment with IFN-γ (fig. S4, B and C). We inferred that DYSF could induce the differentiation and polarization of macrophages.

To elucidate DYSF expression in macrophages, we evaluated its expression in J774A.1 cells that were induced to polarize into defined macrophage phenotypes through exposure to specific cytokines. J774A.1 macrophages were treated with IFN-γ or IL-4, and the expression of traditional macrophage polarization markers was assessed: iNOS and Stat1 for M1-type macrophages, and Arg-1 for M2-type macrophages. Overexpression of DYSF significantly elevated *iNOS* and *Stat1* mRNA levels compared to control cells, indicating that DYSF promotes polarization towards M1-type macrophages as anticipated (Fig. 3C). To further investigate the role of DYSF in M1-type macrophages, we examined the macrophage phenotypes in DYSF siRNA-transfected J774A.1 cells treated with IFN-γ. DYSF-knockdown cells exposed to IFN-γ exhibited markedly reduced *iNOS* and *Stat1* mRNA expression compared to mock and control cells (fig. S4D). Conversely, *Arg-1* mRNA expression was induced in cells treated with IFN-γ and IL-4, but not in DYSF-overexpressing cells (Fig. 3C), suggesting that DYSF has a lesser impact on M2-type macrophage polarization.

**Figure 3.**
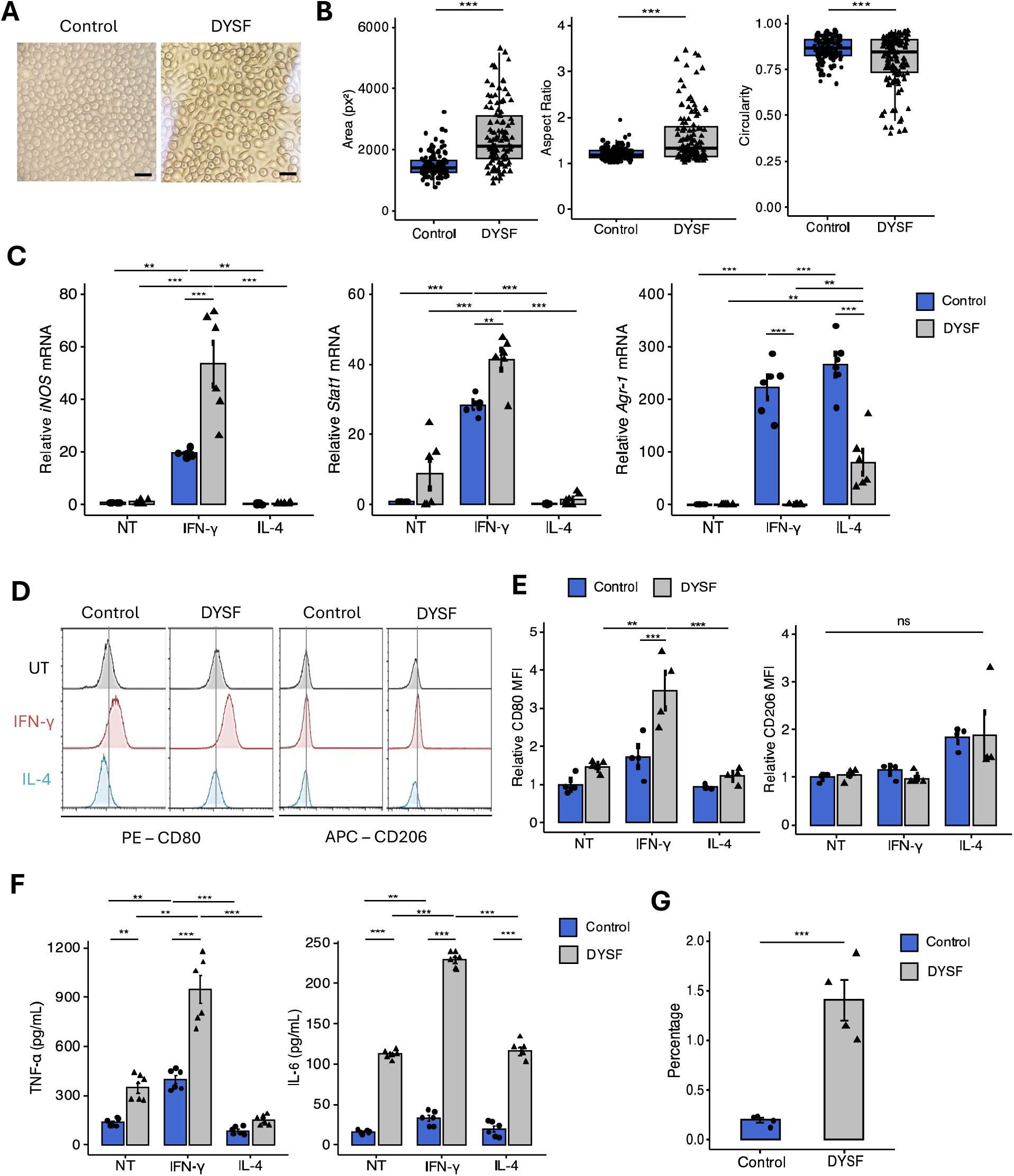
Overexpression of DYSF induces the M1-type macrophages. (A) Representative phase contrast images of J774A.1 transfected with DYSF plasmids (DYSF) or the empty vector control (Control). Scale bar, 50 µm. (B) Quantification of pixel area, circularity and aspect ratio in DYSF or Control vector-transfected J774A.1. One hundred cells were counted for each condition. (C) Expression of *iNOS* (M1), *Stat1* (M1), and *Arg-1* (M2) mRNA was assessed by qPCR to confirm macrophage phenotype. All qPCR reactions were performed in triplicate. (D) Flow cytometry analysis of DYSF or Control vector-transfected J774A.1 treated with 100 ng/mL IFN-γ or 100ng/mL IL-4 for 24 hours. The cells were analyzed according to the expression of Phycoerythrin (PE)-conjugated CD80 (M1) and Allophycocyanin (APC)-conjugated CD206 (M2). (E) Mean fluorescence intensity (MFI) of CD80 (Left) and CD206 (Right) in flow cytometry. n = 4. (F) The protein concentration of TNF-α and IL-6 in conditioned medium obtained from DYSF or Control-transfected J774A.1 was assessed by ELISA. n = 6. (G) The percentages of migration in DYSF or Control vector-transfected J774A.1 to the lower layer of the trans-well assay (n = 4). (B), (C), (E), (F), and (G) were analyzed by unpaired Student’s t-test, and represented as means ± SD. * P < 0.05, ** P < 0.01, and *** P < 0.001. ns, not significant.

To further elucidate the phenotype of macrophage polarization, we conducted flow cytometry analysis based on cell surface marker expression (Fig. 3D). J774A.1 macrophages treated with IFN-γ exhibited elevated levels of CD80, with minimal or absent CD206, resembling M1 macrophages. Overexpression of DYSF in conjunction with IFN-γ treatment resulted in higher CD80 intensity compared to IFN-γ-stimulated control cells, indicating a pronounced tendency towards M1-type macrophage polarization (Fig. 3E). Conversely, cells treated with IL-4 demonstrated increased CD206 expression, suggesting alignment with the M2-type macrophage profile. However, this increase was not statistically significant in either condition, as the mean fluorescence of DYSF-overexpressing cells was similar to that of the control. These findings indicate that DYSF modulates macrophage polarization towards an increased M1-type phenotype.

M1-type macrophages activated by IFN-γ stimulation secrete various pro-inflammatory cytokines. To investigate whether DYSF influences pro-inflammatory macrophage secretion, we analyzed the secretion of TNF-α and IL-6 from the conditioned medium of IFN-γ-stimulated macrophages. DYSF-overexpressing cells, both in the presence and absence of IFN-γ, exhibited higher protein concentrations of TNF-α and IL-6 than control cells (Fig. 3F). Subsequently, we assessed whether the expression of pro-inflammatory cytokines was altered by DYSF knockdown. The ELISA results indicated that DYSF knockdown significantly affected the expression of TNF-α and IL-6 compared to those in the control cells (fig. S4E). These results suggest that DYSF regulates the pro-inflammatory factors, produced by stimulation with IFN-γ, in macrophage.

M1-type macrophages are recognized for their pro-inflammatory function and migrate to sites of infection or injury to initiate an immune response (*17*). To assess the biological function of M1-type macrophages, we conducted transwell assays to determine whether DYSF facilitates cell migration. We cultured DYSF-overexpressing or control J774A.1 cells on Matrigel-coated membranes for 48 hours and subsequently quantified the cells that migrated to the bottom wells across the membrane. DYSF overexpression significantly enhanced macrophage migration to levels comparable to that of the control (Fig. 3G). These findings suggest that DYSF plays a crucial role in macrophage migration.

### DYSF in M1-type macrophages Activates the STAT1 Signaling Pathway

The upregulation of DYSF during macrophage differentiation indicates an associated signaling pathway that may contribute to IFN-γ-induced M1-type macrophage polarization. This polarization is directly linked to STAT1 and is integral to the characteristics of this macrophage subtype (*28*). IFN-γ also enhances the production of TNF-α and IL-6 by activating STAT1, which functions as a transcription factor. To identify the specific molecules activated by DYSF, we investigated the endogenous protein levels and phosphorylation status of key molecules in the STAT pathway. Western blotting revealed that DYSF overexpression affected the levels of unphosphorylated STAT1 (U-STAT1) and the phosphorylation of STAT1 at Ser727 compared to Tyr701 (Fig. 4, A and B). To further explore the involvement of DYSF in the STAT1 signaling pathway, we assessed U-STAT1 and STAT1 phosphorylation in DYSF siRNA-transfected J774A.1 cells. DYSF-knockdown cells exhibited reduced U-STAT1 expression compared to mock and control cells (fig. S5, A and B). Notably, the phosphorylation of STAT1 at Ser727 decreased in DYSF-knockdown cells treated with IFN-γ, whereas it remained unchanged in the other control cells. These findings suggest that the response to DYSF is associated with the phosphorylation of STAT1 at Ser727, implicating its role in inflammatory macrophages.

**Figure 4.**
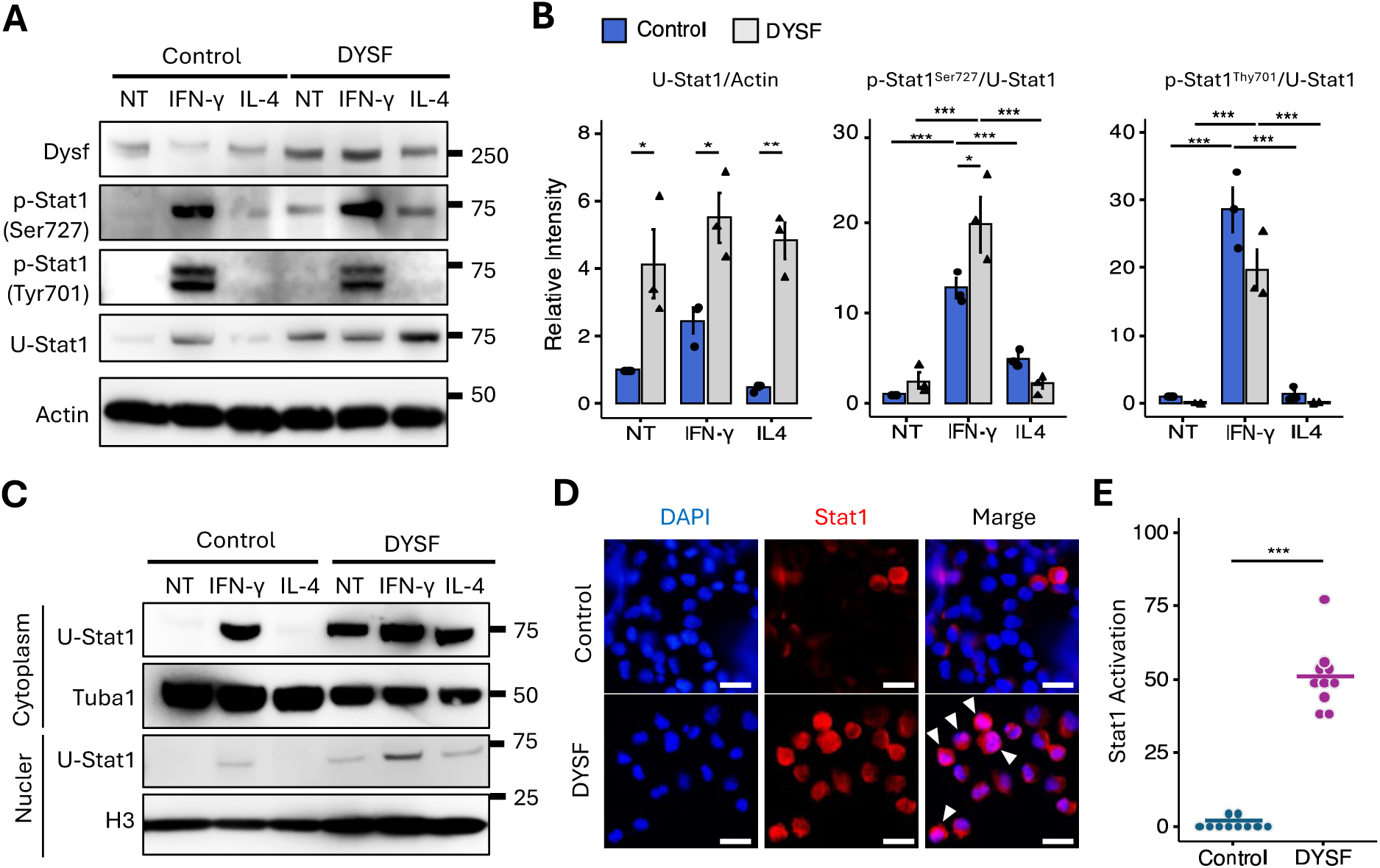
DYSF induces the characteristic of Macrophages via Stat1 pathway. (A) The protein levels of DYSF and the phosphorylation activation of Stat1 in J774A.1 transfected with DYSF plasmids (DYSF) or the empty vector control (Control) were evaluated by western blotting. β-actin (Actin) was used as the loading control. (B) Protein abundance of U-Stat1, Stat1 phospholylated at Ser727 and Tyr701 in J774A.1 treated with or without IFN-γ or IL-4. n = 3. (C) Protein abundance of nuclear and cytoplasm Stat1 in J774A.1 transfected with DYSF or control vector. α-Tubulin (Tuba1) for cytoplasm and Histon H3 (H3) for nuclear were used as the loading control. (D) Representative immunofluorescence images of Stat1 (red) and DAPI (blue), in J774A.1 treated as indicated. White arrow heads indicate the nuclear localization of Stat1. Scale bars, 25 μm. (E) Ratio of nuclear Stat1/DAPI in J774A.1 transfected with DYSF or control vector to assess the Stat1 activation. Cells were counted per field. n = 10. (B) and (D) were analyzed by unpaired Student’s t-test, and represented as means ± SD. * P < 0.05, ** P < 0.01, and *** P < 0.001.

To investigate STAT1 the nuclear translocation of STAT1 in DYSF-overexpressing and control cells, we assessed the amount of U-STAT1 by separating the nuclear and cytoplasmic fractions (Fig. 4C). In DYSF-overexpressing cells, STAT1 was detected in the cytoplasmic extracts regardless of cytokine stimulation when probed with a specific antibody against U-STAT1. Moreover, STAT1 was prominently expressed in the nuclei of DYSF-overexpressing cells upon IFN-γ stimulation, indicating that DYSF plays a role in IFN-γ-induced STAT1 activation. Conversely, the cytoplasmic STAT1 was increased in IFN-γ-stimulated control cells. To verify STAT1 nuclear translocation, we conducted immunofluorescence staining to assess the co-localization of nuclear and STAT1 (Fig. 4D). Immunofluorescence images revealed a clear nuclear accumulation of U-STAT1 in DYSF-overexpressing cells compared to control cells (Fig. 4E). These findings demonstrate that DYSF is a critical regulator of the STAT1 signaling pathway, facilitating not only the phosphorylation of STAT1 at Ser727 but also its nuclear translocation of U-STAT1.

### DYSF Induces STAT1 Activation via Ca^2+^ Influx in Macrophages

Calcium influx can modulate STAT1 activity, including phosphorylation and subsequent nuclear translocation, while STAT1 itself can influence calcium signaling pathways and related gene expression (*29*). To investigate the effect of DYSF on calcium signaling within the STAT1 signaling pathway in macrophages, we first monitored intracellular Ca^2+^ levels in J774A.1 cells under conditions of both DYSF overexpression and knockout. Addition of A23187 (2 µM), a Ca²⁺ ionophore increased the Fura-2 ratio in presence of calcium, indicating calcium bounding within macrophages (fig. S6). DYSF-overexpressing cells exhibited a significant increase in intracellular Ca^2+^ following treatment with A23187 compared to control cells (Fig. 5, A and B). Furthermore, IFN-γ treatment of DYSF-overexpressing cells resulted in a subsequent increase in intracellular Ca^2+^ levels, suggesting that DYSF can mediate intracellular Ca^2+^ influx in conjunction with the IFN-γ signaling pathway. Conversely, when we monitored calcium influx in J774A.1 cells with reduced DYSF expression, the calcium influx following A23187 treatment was significantly diminished in DYSF-knockout cells compared to that in mock and control siRNA-transfected cells (Fig. 5, C and D). To assess the influence of intracellular Ca^2+^ on the STAT1 signaling pathway in macrophages, we treated J774A.1 macrophages expressing DYSF or control cells with A23187. The abundance of U-STAT1 and phosphorylated STAT1 in DYSF-overexpressing cells increased with A23187 treatment, whereas the control cells were minimally affected (Fig. 5, E and F). These results suggested that the intracellular Ca^2+^ affects STAT1 protein levels in a manner dependent on DYSF expression. We also assessed the localization of U-STAT1 in DYSF-expressing cells treated with A23187. Immunofluorescence images revealed that treatment of J774A.1 macrophages with A23187 increased nuclear localization of STAT1 (Fig. 5, G and H), indicating that intracellular Ca^2+^ contributes to the activation of STAT1 in the nucleus of macrophages.

**Figure 5.**
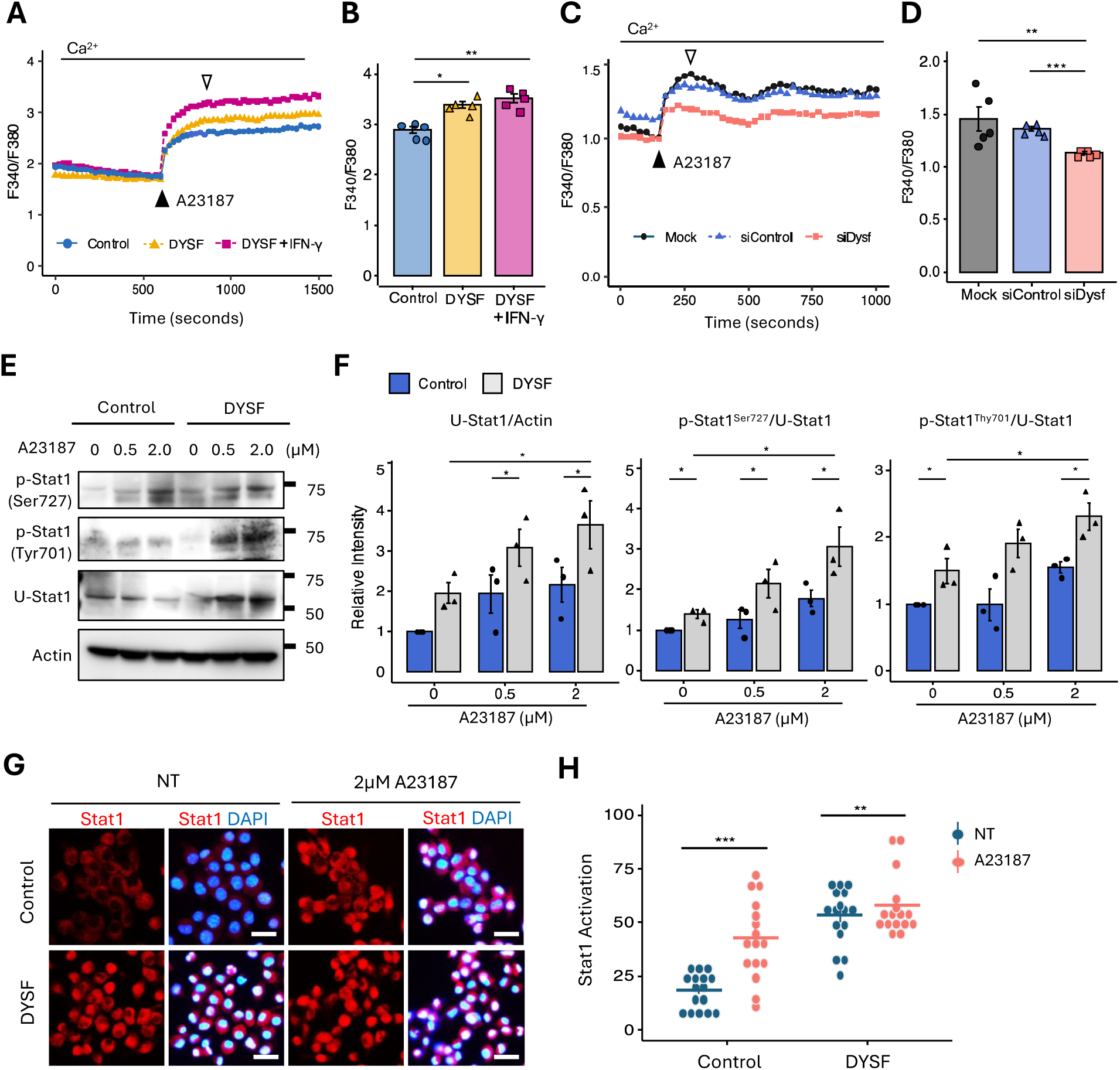
DYSF-Mediated Calcium Influx Regulates STAT1 in Macrophages. (A) Time-course of fluorescent signal for intracellular calcium concentration before and after application of 2 µM A23187, a Ca²⁺ ionophore in J774A.1 transfected with DYSF plasmids (DYSF) or the empty vector control (Control). J774A.1 transfectd with DYSF plasmids were treated with or without 100 ng/mL IFN-γ for 24 hours. The ratio of fluorescence was determined by changes at 340 and 380 nm (F340/380). (B) Summary of the maximal luminescence signals (white arrow heads) measured after application of ionomycin in J774A.1 transfected with DYSF or Control vector. n = 5. (C) Time-course of fluorescent signal for intracellular calcium concentration before and after application of 2 µM A23187 in J774A.1 transfected with DYSF siRNA (siDYSF) or non-targeting siRNA (siControl), and non-transfected cells (Mock). Calcium concentration was measured by fluorescence of Fluo-2. The ratio of fluorescence was determined by changes at 340 and 380 nm (F340/F380). (D) Summary of the maximal luminescence signals (white arrow heads) measured after application of ionomycin in J774A.1 transfected with siDYSF or siControl, and Mock. n = 5. (E) The protein levels of the phosphorylation activation of Stat1 in DYSF or Control vector-transfected J774A.1 treated with or without A23187 were evaluated by western blotting. β-actin (Actin) was used as the loading control. (F) Protein abundance of U-Stat1, Stat1 phospholylated at Ser727 and Tyr701 in J774A.1 treated with or without A23187 (n = 3). Density of the protein bands was normalized to Actin or Stat1. (G) Representative immunofluorescence images of Stat1 (red) and DAPI (blue), in J774A.1 transfected as indicated. Scale bars, 25 μm. (H) Ratio of nuclear Stat1/DAPI in J774A.1 transfected with DYSF or control vector. Cells were counted per field (n = 16). (B) and (D) were analyzed by One-wayANOVA and Tukey-Kramer test. (F) and (H) were analyzed by unpaired Student’s t-test. Figures [(B), (D), (F), and (H)] represented as means ± SD. * P < 0.05, ** P < 0.01, and *** P < 0.001.

### DYSF in Macrophages Alters Skeletal Muscle Fiber Fusion

Macrophages are integral to the fusion of skeletal muscle fibers, a process vital for muscle regeneration and repair (*30*). M1-type macrophages are involved in regulating inflammation, phagocytosis, and the recruitment and activation of muscle stem cells (satellite cells, MuSCs). To elucidate the impact of DYSF-regulated differentiation of M1 macrophages on muscle fiber formation, we co-cultured mouse monocyte/macrophage J774A.1 cells with mouse myoblast C2C12 (Fig. 6A). DYSF-overexpressing and control vector-transfected macrophages were cultured in the upper well of a Boyden chamber insert, while C2C12 cells were seeded in the lower well. For myoblast differentiation and fusion, cells were cultured in a medium containing 2% horse serum, and myoblast differentiation into myotubes was monitored over time. Myofiber fusion was assessed by western blotting and immunohistochemistry using muscle differentiation markers, myosin heavy chain 1 (Myh1), Dysf, and Myogenin (Myog). In the control co-culture, differentiation induction led to increased expression of Dysf and Myog, whereas the expression of these proteins was suppressed in DYSF-overexpressing macrophages (Fig. 6, B and C). Furthermore, immunofluoresence staining for DYSF and Myh1 demonstrated that differentiation induction increased both expression and enhanced fusion into myotubes in the control co-culture, whereas fusion was suppressed in the DYSF-overexpressing monocyte co-culture (Fig. 6, D and E). Additionally, the fusion index of Myh1-positive myotubes was measured, revealing that myoblasts co-cultured with DYSF-overexpressing macrophages exhibited a lower fusion index than those in the control (Fig. 6F). These finding that DYSF is a key factor for altering the properties of macrophages, leading to myoblast fusion (Fig. 6G).

**Figure 6.**
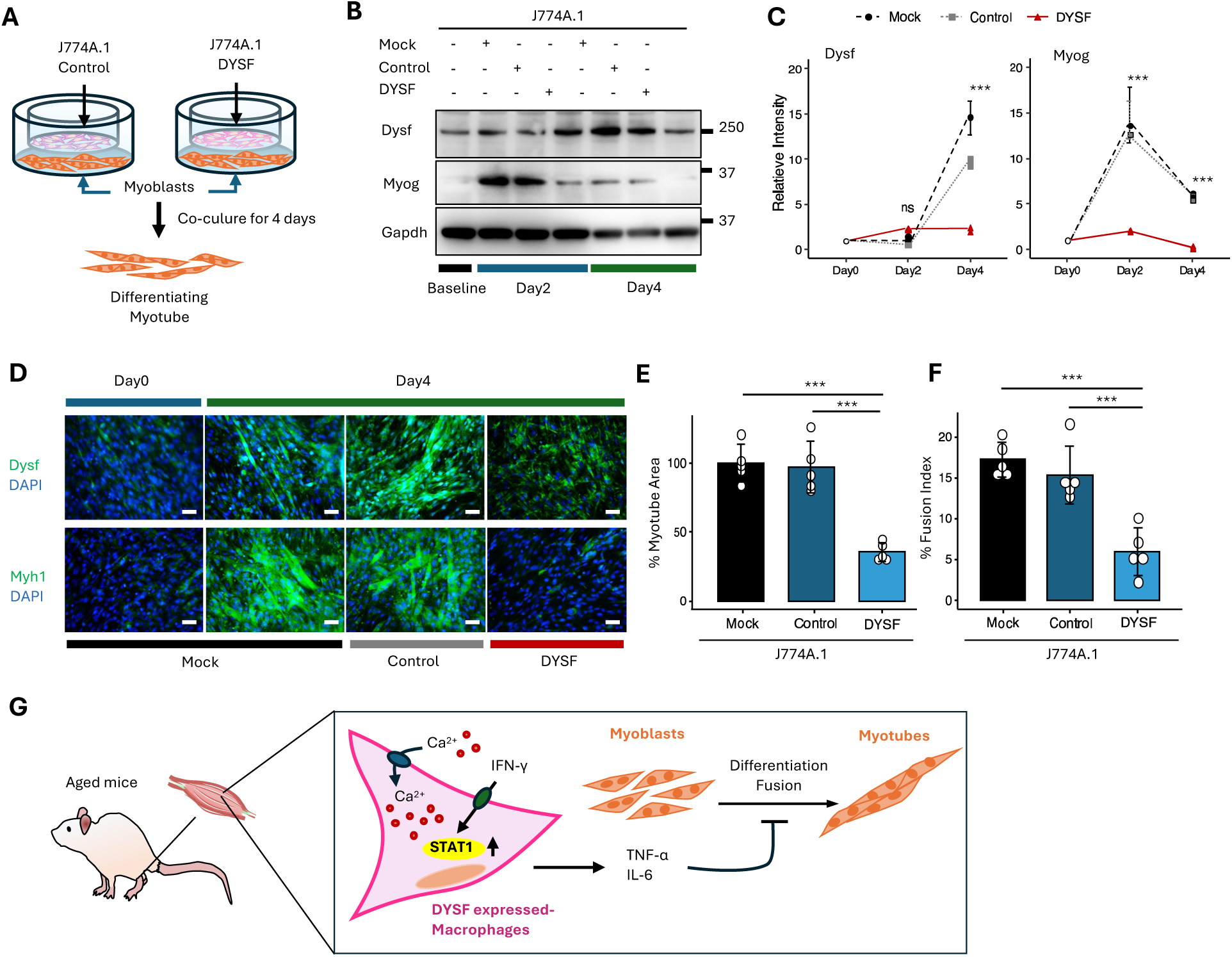
DYSF-Driven Macrophage Changes Suppress Myotube Fusion. (A) Schematic illustration of co-culture system using in the experiments with J774A.1 macrophages and C2C12 myoblasts. (B) Western blot analysis of differentiated C2C12 cell lysetes after co-cultured with J774A.1 DYSF plasmids (DYSF) or the empty vector control (Control), and non-transfected cells (Mock) with antibodies against DYSF and Myogenin (Myog), a transcriptional activator for myogenesis. GAPDH used as the loading control. (C) Relative intensity of protein abundance of DYSF in differentiated C2C12 after co-culture J774A.1 transfected with DYSF or control vector, and Mock. n = 3. (D) Representative immunofluorescence images of Dysf (green, upper) and Myh1 (green, lower), in C2C12 after co-cultured with J774A.1 transfected with DYSF or Control vector, and Mock. Scale bars, 50 μm. (E) Quantitative intensity for Myh1-positive myotubes was performed four days after myoblast differentiation. Myh1-positive myotubes were assessed per field. n = 5. (F) Quantitative analysis for the number of Myh1-positive myotubes with more than 5 nuclei was performed four days after myoblast differentiation. Fusion index was calculated as the ratio of nuclei in myotubes to the total number of nuclei. n = 5. (G) Model showing the positive regulation of IFN-γ/STAT1 pathway though the accumulation of intracellular Ca^2+^ in DYSF-overexpressing macrophages. Activated DYSF-overexpressing macrophages can release pro-inflammatory cytokines, which inhibit the differentiation of myoblasts. (C), (E), and (F) were analyzed by One-way ANOVA and Tukey-Kramer test, and represented as means ± SD. *** P < 0.001. ns, not significant.

## Discussion

Although DYSF is highly expressed in both monocytes and myofibers, its role in monocyte differentiation is unclear. In this study, we report that DYSF expression increases with age in monocytes, preferentially inducing differentiation into M1-type inflammatory macrophages within the skeletal muscle. M1-type macrophages activated by DYSF overexpression engage the STAT1 signaling pathway via Ca²⁺ influx, resulting in TNF-α and IL-6 secretion. Furthermore, DYSF-induced M1-type macrophages suppress muscle cell fusion through immune-skeletal muscle interactions. To our knowledge, this is the first report linking the hyperaccumulation of DYSF in monocytes to the loss of muscle differentiation with age.

*Dysf*-deficient muscle exhibits persistent accumulation of macrophages, which transition to a pro-inflammatory, cytotoxic phenotype, exacerbating muscle fiber damage. This suggests that dysferlinopathy is not solely a repair defect but also involves an immune-mediated degenerative process. Mild myofiber damage in *Dysf*-deficient muscles initiates an inflammatory cascade, underpinning a myofiber-specific dystrophic process. Monocytes derived from *Dysf*-deficient SJL mice and patients with dysferlinopathy demonstrate increased phagocytic activity towards damaged myofibers (*14*). In response to their endocytic activity, Rho family small GTPases are upregulated, regulating phagocytosis to rescue *Dysf*-deficient myofibers, resulting in a slowly progressive, late-onset disease. DYSF modifies the capacity of monocytes to perform these functions via the small Rho family of GTPases. Furthermore, *Dysf*-deficient muscles show chronically elevated IL-1β levels (*24*). Increased IL-1β impairs muscle regeneration by disrupting the transition from M1 to M2-type macrophages. Inhibition of IL-1β signaling improved regeneration, indicating its potential as a therapeutic target. Mononuclear cells directly influence myofiber regeneration in *Dysf*-deficient muscle. Transplantation of bone marrow from wild-type mice into *Dysf*-deficient BLA/J mice restored muscle function (*16*). However, the presence of anti-inflammatory monocytes/macrophages expressing CD206 and Arg-1 did not significantly increase in either group, failing to prevent disease progression. These findings suggest that loss of DYSF expression in muscles promotes intramuscular expansion of inflammatory macrophages. Nonetheless, the impact of DYSF expression in macrophages on myogenic differentiation remains unclear. In contrast, our results demonstrated that the stability of DYSF in monocytes/macrophages increased migration and inflammation signaling, indicating that DYSF contributes to macrophage activation. According to data from other groups, DYSF is overexpressed in monocytes from patients with atherosclerotic cardiovascular disease (ASCVD) due to hypermethylation of the DYSF promoter, suggesting that DYSF enhances monocyte activation, including adhesion, migration, and phagocytosis (*13*). Hypermethylation of the DYSF promoter leads to the expression of Selectin L, an adhesion molecule that facilitates interactions between monocytes and vascular endothelial cells. Although the underlying reasons for the discrepancies between these two studies remain unclear, the functions of DYSF may vary depending on the specific factors activated within monocytes/macrophages. Our findings indicate that DYSF plays a central role in M1/M2 macrophage polarization via Ca²⁺ signaling cascades. The influx of Ca²⁺ is crucial for macrophage polarization, as elevated intracellular Ca²⁺ levels regulate cytokine production and support the associated inflammatory signaling pathways (*31*). Furthermore, the accumulation of cytosolic Ca²⁺ signals in aging mice leads to the activation of the NF-κB signaling pathway and the production of pro-inflammatory cytokines, TNF-α, and IL-6 (*32*). The regulation of Ca²⁺ concentration by DYSF may determine whether macrophages promote inflammation or facilitate repair, and these findings may serve as a foundational contribution to understanding macrophage polarization.

The transition between M1 and M2 macrophage phenotypes is intricate. The equilibrium between the activation of STAT1 and STAT3/STAT6 plays a crucial role in regulating macrophage polarization and activity (*33*). The predominant activation of NF-κB and STAT1 facilitates M1 macrophage polarization, leading to cytotoxic and inflammatory functions. In this study, we demonstrated that DYSF positively influences STAT1 stability and function via Ca²⁺ influx. STAT1 is primarily activated by interferons, such as IFN-α/β and IFN-γ, as well as IL-6 (*34*). Upon activation, JAK kinases phosphorylate STAT1, which then dimerizes, translocates to the nucleus, and regulates transcription. Ca²⁺ influx may indirectly modulate STAT1 activity. Calcium–calmodulin (CaM)-dependent protein kinase II (CaMKII) and Ca²⁺-dependent kinases enhance STAT1 phosphorylation (*29*). Consequently, intracellular Ca²⁺ serves as a fine-tuner of STAT1 the strength and duration. In macrophages, STAT1 signaling promotes the expression of pro-inflammatory cytokines that affect store-operated Ca²⁺ entry (SOCE) and ER–mitochondria Ca²⁺ transfer, suggesting the promotion of M1 polarization (*31*). Our findings indicate that DYSF contributes more significantly to the stability and nuclear translocation of STAT1 than to its phosphorylation in macrophages than to its phosphorylation. Additionally, Ca²⁺ signaling may directly regulate STAT1 rather than through Ca²⁺-dependent kinases and phosphatases.

Skeletal muscle mass, strength, and regenerative capacity decline with age. Aging also impedes the regeneration and repair of skeletal muscle following injury (*35*). These processes are characterized by a reduction in muscle stem cells (MuSCs), impaired muscle fiber function, and accumulation of adipose and connective tissues. Additionally, there is selective atrophy of fast-twitch Type II fibers, a decrease in motor units, and alterations in the cellular and extracellular environments, including changes in fibro-adipogenic progenitors (FAPs) and the extracellular matrix. These cellular and molecular changes culminate in diminished muscle quality, reduced elasticity, and compromised ability to repair and function effectively. Cytokines secreted by macrophages are crucial for MuSC stimulation and play a significant role in muscle regeneration after injury. Following injury, M1-type macrophages initially infiltrate the damaged area, clearing necrotic debris and releasing cytokines such as TNF-α, IL-1, IL-6, and chemokines that promote MuSC activation and proliferation (*36*, *37*). Subsequently, macrophages transition to an anti-inflammatory, pro-regenerative phenotype characterized by the production of IL-10, Insulin-like Growth Factor (IGF)-1, and Transforming growth factor-β (TGF-β), which facilitate MuSC differentiation and tissue remodeling. This dynamic shift from M1 to M2 is essential for effective tissue regeneration. However, during aging, the recruitment and polarization of macrophages become dysregulated. Aged macrophages frequently display a persistent pro-inflammatory phenotype, marked by increased secretion of TNF-α and IL-1β and diminished production of pro-regenerative factors. This chronic low-grade inflammation disrupts the timely M1-to-M2 transition, thereby impairing the regenerative milieu. Consequently, MuSCs experience prolonged inflammatory stress, resulting in impaired proliferation, exhaustion, and inappropriate differentiation. Our findings indicate that DYSF expression increases with age, leading to the accumulation of M1-type macrophages. Overexpression of DYSF in macrophages inhibits myoblast differentiation and fusion, underscoring the critical role of DYSF in maintaining myogenesis in both skeletal muscles and macrophages. DYSF-overexpressing macrophages secrete TNF-α and IL-6, suggesting that pro-inflammatory cytokines may be constitutively present in the skeletal muscle niche, stimulating MuSCs. Thus, DYSF in macrophages may be a pivotal factor in maintaining MuSCs.

In our study, DYSF emerged as a significant factor in the interaction between the immune system and skeletal muscle during aging. The observed increase in DYSF expression in monocytes with advancing age suggests that DYSF may perform a function distinct from its role in dysferlinopathy. Consequently, it is imperative to elucidate the mechanisms underlying the age-related elevation of DYSF protein levels. Previous research has indicated that the DNA methyltransferase DNMT1 modulates DYSF promoter methylation in ASCVD (*13*), implying that the methylation patterns of DYSF or age-related genes may be altered with aging. Additionally, it is crucial to consider that the accumulation of DYSF proteins with age may be linked to the mechanisms of protein aggregation and degradation, such as endoplasmic reticulum (ER) stress and autophagy. This study suggests that the DYSF-related Ca²⁺/STAT1 signaling pathway may serve as a potential therapeutic target for sarcopenia, a condition characterized by age-related muscle atrophy. Given that Ca²⁺ flux is essential for DYSF-induced macrophage polarization, the development of small-molecule inhibitors targeting Ca²⁺ is significant for aging-suppressive effect. Furthermore, assessing Ca²⁺ concentration or DYSF protein levels in blood samples from aging individuals may prove valuable. This hypothesis should be validated by examining a larger cohort of older adults in future studies.

## Materials and Methods

### Animals

Mouse experiments were performed under protocols approved by the Yamaguchi University Animal Care and Use Committee (No.80-002 and No.80-021). Male and female were used in this study, and sex-specific features of age were analyzed independently. Mice were group-housed in ventilated cages with a constant temperature of 25 °C, 30–70% humidity and a 12-h light/dark cycle. Water and the PicoLab Rodent Diet 20 (LaboDiet) were provided ad libitum.

At the time of bone marrow collection, young mice were 6–8 months of age and old mice were 18– 30 months of age. After euthanasia, bone marrow samples from limbs were centrifuged (10,000 g) for 30 seconds at 4 °C and collected bone marrow cells according to a protocol (*38*). To isolate mouse monocyte, cells obtained from bone marrow samples were incubated with a mixture of Monocyte Biotin-Antibody Cocktail (Miltenyi Biotec) and separated using the MACS system according to the manufacturer’s instructions (Miltenyi Biotec).

Isolated monocytes were cultured in high glucose Dulbecco’s Modified Eagle Medium (DMEM; Nacalai Tesque) containing 10 % fetal bovine serum (FBS, Nichirei), 200 mM GlutaMAX (Invitrogen), 5 ng/mL recombinant mouse GM-CSF (Biolegend), 2.5 ng/mL recombinant mouse IL-4 (Biolegend), 100 units/ml penicillin (Nacalai Tesque), and 100 µg/ml streptomycin (Nacalai Tesque) at a non-treated surface plate in a humidified atmosphere at 37°C in 5% CO_2_. The culture medium was changed every 2 days. To isolated monocyte-derived CD11b+ macrophages, cultured cells were incubated with a biotin-conjugated antibody against CD11b (BD Biosciences) for 30 min at 4°C and isolated by BD IMag Cell Separation System according to the manufacturer’s instructions.

### Cell Culture

J774A.1, U937 and C2C12 were purchased from the American Type Culture Collection. J774A.1, a murine monocyte/macrophage cell line was cultured in high glucose DMEM containing 10 % FBS, 100 units/ml penicillin, and 100 µg/ml. Cell lines were cultured in a humidified atmosphere at 37 °C in 5 % CO_2_, and the culture medium was changed every 2 days. When cells reached subconfluence, they were rinsed in phosphate-buffered saline (PBS, pH 7.4, Nacalai Tesque) and dislodged the cells using a cell scraper for passage.

U937 cells, a human pro-monocytic model cell line was cultured in a T75 flask in RPMI 1640 medium without phenol red (RPMI; Nacalai Tesque) containing 10 % fetal bovine serum (FBS, Nichirei), 100 units/ml penicillin (Nacalai Tesque), and 100 µg/ml streptomycin (Nacalai Tesque). Cells were passaged for a maximum of 4 weeks. For differentiation to macrophage, cells (1 x 10^6^ cells) were passaged to a 6-well plate and treated with 10 nM phorbol-12-myris-tate-13-acetate (PMA, FUJIFILM Wako Pure Chemical Corp.) for 24 or 48 hours.

C2C12, a mouse myoblast cell line was cultured in high glucose DMEM containing 10 % FBS, 100 units/ml penicillin and 100 µg/ml streptomycin. For studies of myogenic differentiation, when cells were reached at full confluency, the culture medium was substituted with differentiated medium contained with DMEM with 2% horse serum (Thermo Fisher Scientific). Differentiated medium was changed every day.

### Cell proliferation assay

J774A.1 cells were seeded in 96-well plates at 10,000 cells/well. After incubated for 24, 48, and 72 hours, cell proliferation was measured using Cell Titer 96 AQueous Assay Kit (Promega) according to the manufacturer’s specifications.

### Small interfering RNA (siRNA)

For knockdown of DYSF, we used Silencer Select Mouse DYSF (Assay ID s77352) purchased from Invitrogen. J774A.1 cells were seeded at a density of 1 × 10^5^ cells/well into a 6-well plate and incubated with siRNAs (15nM) and Lipofectamine RNAiMAX transfection reagent (Invitrogen) according to the manufacturer’s protocol. The control was Silencer Select Negative Control No. 1 siRNA. siRNA-transfected cells were incubated at 37°C in 5% CO_2_ until the assay. Representative data from three independent experiments.

### Transient transfection with plasmids

DYSF-3HA was a gift from Steven Vogel (Addgene plasmid # 29767; http://n2t.net/addgene:29767; RRID: Addgene_29767) (*39*). The control was a pcDNA3.1 vector. For transient transfection of plasmids, J774A.1 cells were seeded at a density of 6 × 10^5^ cells per well into a 6-well plate. After incubation for 24 hours, the plasmids (10 μg) and a Polyethylenimine MAX (PEI MAX, Polysciences) in culture medium were added to each well. After 4 hours incubation, we changed culture medium and incubated until the experiment is performed.

### In vitro migration assay

1 × 10^5^ of plasmid-transfected J774A.1were seeded on Matrigel (Corning)-coated insert (pore size, 5.0 µm) in serum-free medium. After 24 hours, the cells attached to the lower side were counted. The invading cells in four randomly fields were counted, and the mean of counted cell were reported.

### Intracellular calcium level observation

The Ca²⁺ levels were determined with the use of Calcium Kit II-Fluo 2 (Dojindo). Control vector or DYSF vector-transfected J774A.1 cells were plated at 4 x 10^4^ cells per well into the 96-well plates for 24 hours. After incubation, cells were incubated with 4 μg/ mL Fluo 2-AM with 5% Pluronic F-127 and 250 mmol/L Probenecid at 37 °C in 5 % CO_2_ for 1 hour. To evaluate the internal stores of calcium, we added 0.5 or 2 µM A23187, a calcium ionophore during the measurement. Cells measurement was performed on a Flexstaion 3 (Molecular Devices, USA) with an excitation wavelength of 340/510 nm and an emission wavelength of 380/510 nm. Baseline signals (F0) were recorded 5 min before the addition of each stimulus. Subsequently, continuous fluorescence measurements were performed for 20 min each 25 seconds. Results are shown as F/F0 ratios after background subtraction, where F was the fluorescence signal intensity and F0 was the baseline intensity, as calculated by the average of 5 frames before stimulus application.

### Nuclear and cytoplasmic protein extraction

To separate nuclear and cytoplasmic protein extraction from cells, the cell fractionation was done following previously protocol with modifications (*40*). Breifly, J774A.1 cells (1×10^6^) were collected and washed in PBS. To collect the cytosolic fraction, the pellets were resuspended in cold lysis buffer containing with 10mM HEPES, 1.5mM MgCl2, 10 mM KCl, 0.1mM EDTA, 0.1% NP-40, a phosphatase inhibitor cocktail (FUJIFILM Wako Pure Chemical Corp.) and a protease inhibitor (Roche). After centrifugation, the nuclear pellets were resuspended in cold buffer containing with 20mM HEPES, 1.5mM MgCl_2_, 400nM NaCl, 0.1mM EDTA and 10% Glycerol and homogenate with the 27G needle. After centrifugation, we collected the supernatant as the nuclear extract.

### Western blotting

The cultured cells in 6-well plates were washed with PBS and then lysed with lysis buffer containing M-PER Mammalian Protein Extraction Reagent (Thermo Scientific) with a phosphatase inhibitor cocktail and a protease inhibitor. The proteins were separated by 10 % SDS-PAGE and transferred to polyvinylidene fluoride (PVDF) membranes. The membranes were blocked with Block Ace (non-fat skim milk, BioRad) in distilled water at room temperature for 30 min. The PVDF membranes were incubated with primary antibodies (table S1) at room temperature for 1 hour. After washing three times in PBS containing 0.1% Tween-20, the PVDF membranes were incubated with horseradish peroxidase (HRP)-conjugated anti-rabbit IgG (Cell Signaling Technology) or anti-mouse IgG (GE Healthcare Life Science) as secondary antibodies at room temperature for 1 hour. Signals were developed using an Immobilon Western Chemiluminescent HRP Substrate (Cytiva) and examined using an Amasham imaging system (GE Healthcare Life Science). To analyze the intensity of bands, we used Fiji.

### Histochemical and Immunofluorescence staining

Frozen 8-µm thin sections of the mouse soleus muscle or cultured cells were fixed 4% paraformaldehyde/PBS. For histochemical staining, sections were stained by Hematoxylin and Eosin Stain Kit (Vector Laboatories). For immunofluoresence staining, sections or cells washed with PBS containing 0.1% Triton X-100 to permeabilize samples and incubated with 3 % bovine serum albumin (BSA, Nacalai Tesque) at room temperature for 60 min to block non-specific binding. Samples then incubated at 4 °C overnight with the primary antibodies (table S1). After washing with PBS, the cells and sections were incubated with an Alexa Fluor 488 Goat Anti-Rabbit IgG H&L (ab150077, abcam), Alexa Fluor 488 Goat Anti-Mouse IgG H&L (ab150113, abcam), Alexa Fluor 594 Goat Anti-Rabbit IgG H&L (ab150080, abcam), or an Alexa 594 anti-Mouse IgG antibody (ab150116, abcam) as a secondary antibody at room temperature for 1 hour. the cells and sections were enclosed using ProLong Glass Antifade Mountant (Thermo Fisher Scientific) after staining the nuclei with 4’,6-diamidino-2-phenylindole (DAPI) at room temperature for 5 min. The fluorescent images were obtained under a fluorescence microscope (Keyence). The total number of signal dots was counted separately in the nucleus and the cytoplasm using Fiji.

### Flow cytometry for the detection of proteins located on the extracellular surface of cells

Collected cells (1 x 10^6^ cells) incubated with PE anti-mouse CD80 Antibody (104707, Biolegend) or APC anti-mouse CD206 (MRC1, MMR) Antibody (141707, Biolegend) that diluted with DMEM containing 2 % FBS at 4 °C for 30 min. For quality control tested, we used PE Mouse IgG2a, κ Isotype Ctrl (FC) Antibody (400213, Biolegend) or APC Rat IgG2a, κ isotype control (400511, Biolegend) under the same conditions. After washing with PBS, cells were stained with DAPI (D523, Dojindo) to exclude dead cells. All flow cytometric analyses were performed with a NovoCyte Flow Cytometer (Agilent). Obtained data were analyzed using the FlowJo software (Treestar).

### Enzyme-linked immuno-sorbent assay (ELISA)

Concentrations of the pro-inflammatory cytokines, TNF-α and IL-6 were determined from J774A.1 cell (1 x 10^6^ cells) culture supernatants using ELISA MAX Standard Set Mouse TNF-α (430901, BioLegend), ELISA MAX Standard Set Mouse IL-6 (431301, BioLegend) following the manufacturers’ instructions. We measured absorbance at 450 nm using a Flexstaion 3 (Molecular Devices) within 30 min after adding TMB Substrate.

### Quantitative RT-PCR (qPCR)

Total RNA was prepared using the ISOSPIN Cell & Tissue RNA (Nippon Gene) and was transcribed into cDNA using the ReverTra Ace qPCR RT Master Mix with gRNA remover (Toyobo). The qPCR was performed using PowerTrack SYBR Green Master Mix for qPCR (#A46109, Thermo Fisher Scientific) with specific forward and reverse primers (table S2) in a QuantStudio3 (Thermo Fisher Scientific). The relative gene expression was calculated with the ΔΔCt method (Livak and Schmittgen, 2001), using GAPDH for normalization.

### Statistics

Statistical analysis and visualized data were performed using R software (v4.5.1). We used unpaired Student’s t-test to compare differences between two samples. We also used one-way ANOVA and Tukey-Kramer test for differences between the variances of the three groups. Values are presented as means ± S.D.

## Acknowledgments

We thank A. Kobayashi for supporting general lab work. We thank Science Research Center, Insutitute of Life Science and Medicine for animal research, and the Yamaguchi University Center for Gene Research for using Flow cytometry, Imaging system, Microplate reader, and Real-time PCR system.

## Founding

This work was supported by Astellas Foundation for Research on Metabolic Disorders (K.T.), Uehara Memorial Foundation (K.T.), New Frontier Research Grant by Yamaguchi University (K.T., N.T.), Finding-Out & Crystallization of Subliminals (FOCS) project by Yamaguchi University of Medicine (K.T., N.T.). This work was supported in part by a Grant-in-Aid for Research Activity Start-up (21K21219), Early-Career Scientists (22K17826) from Japan Society of the Promotion of Science (JSPS) (N.T.), ACT-X (JPMJAX222B) from Japan Science and Technology Agency (JST) (N.T.), the Ube City Next-Generation Researchers Project (N.T.). This work was the result of using research equipment shared in MEXT Project for promoting public utilization of advanced research infrastructure (Program for supporting construction of core facilities) Grant Number JPMXS0440400023.

## Author contributions

K.T. conceived and designed the study. K.T. and N.T. performed experiments and did the data analysis and interpretation. N.T. provided the tissue preparation. K.T. wrote the manuscript and prepared the figures. K.T. and N.T. reviewed and edited the paper.

## Data and materials availability

All data in this published article (and its Supplementary Information files) are available. The other data analyzed in the current study are not publicly available for ethical reasons.

