## Supplementary Files for "Age-dependent Dysferlin Accumulation in Macrophages Promotes STAT1 Activation via Calcium Influx, Impairing Myogenesis"

### Supplementary Figures

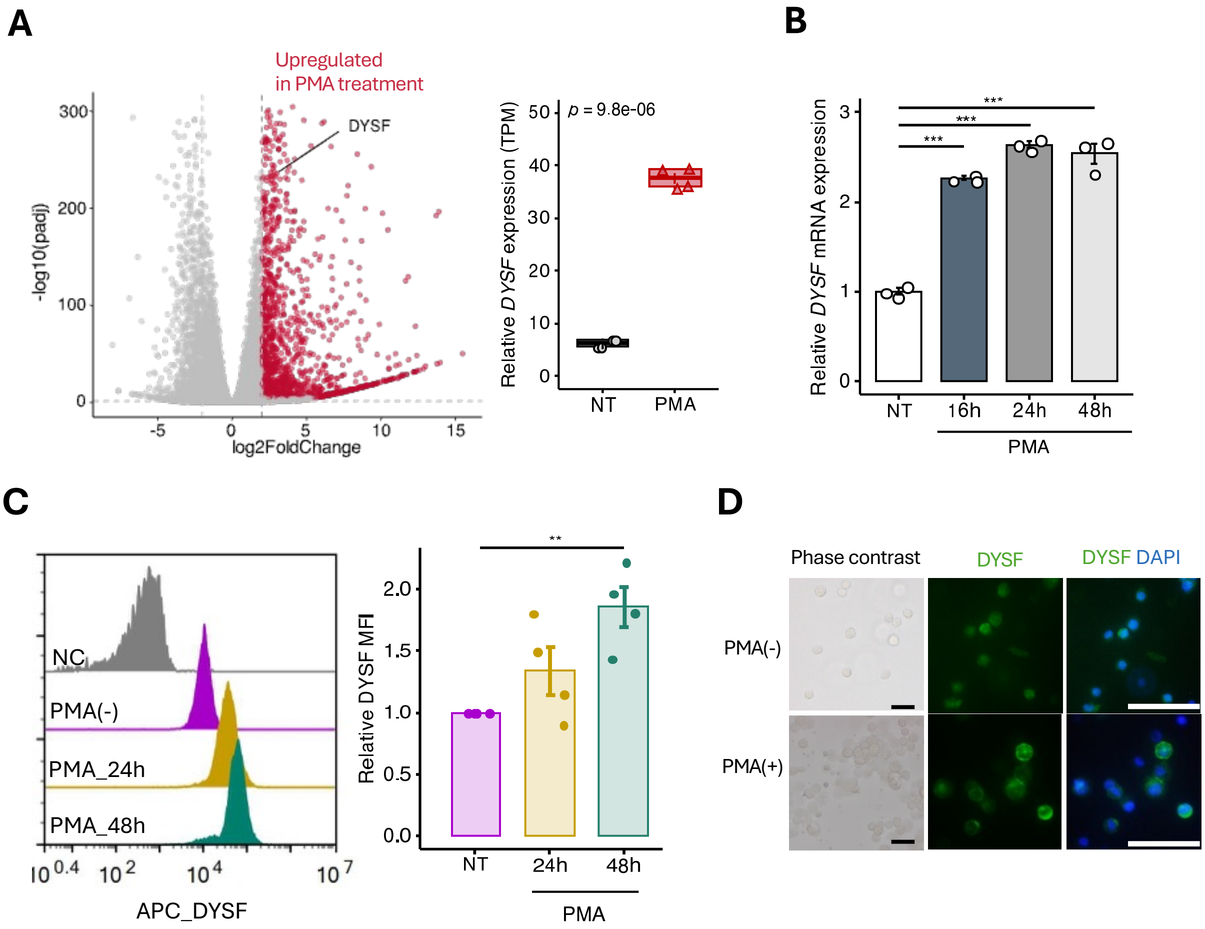


**Fig. S1 DYSF is upregulated in PMA-induced Monocyte Differentiation.**

(A) Volcano plot of the transcriptional microarray data (GSE107566) with or without phorbol 10 nM 12-myristate 13-acetate (PMA) in U937 (left). The transcriptional levels of DYSF were log-normalized by the $\log_2$(TPM) method (right). NT, Non-treated. (B) Expression of *DYSF* mRNA was assessed by qPCR in U397 treated with PMA in time-dependent manner. n = 3. (C) Flow cytometry analysis of U937 treated with PMA for 24 or 48 hours using an antibody against DYSF (left). NC; negative control. Mean fluorescence intensity (MFI) of DYSF expression in flow cytometry (right). n = 4. (D) Representative phase contrast and immunofluorescence staining images with an antibody against DYSF in U937 treated with or without PMA. DAPI was used as a nuclear stain. Scale bars, 25 µm. (A) (right) was analyzed by unpaired Student's t-test. (B) and (C)(right) were analyzed by One-way ANOVA and Tukey-Kramer test. Figures [(A), (B), and (C)(right)] represented as means ± SD. ** $P < 0.01$, and *** $P < 0.001$.

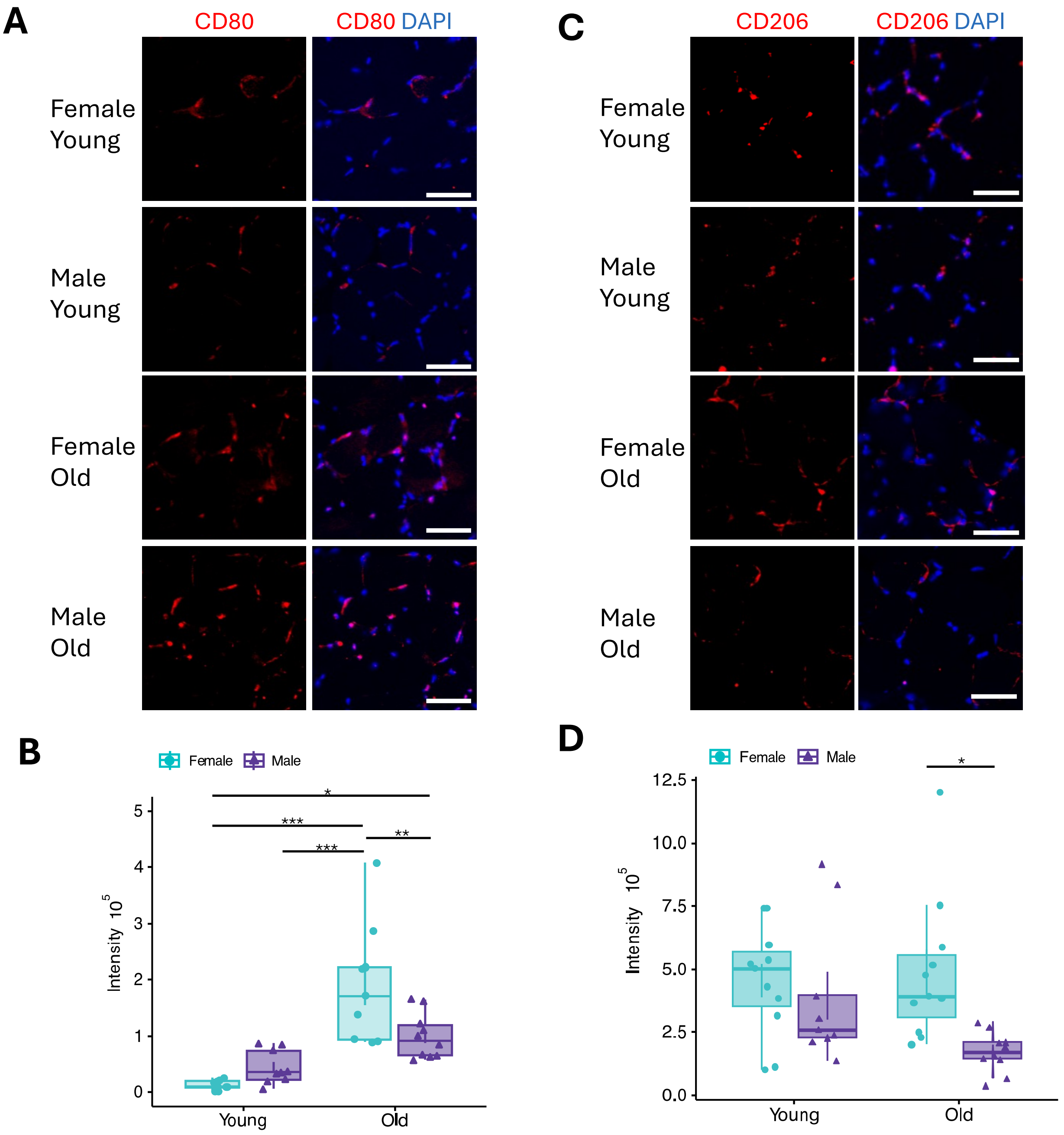


**Fig. S2 Pro-Inflammatory Macrophage Infiltration in Aged Muscle.**

(A) Representative images of frozon sections of soleus muscle, stained with antibodies against CD80 for M1-type macrophages. DAPI was used as a nuclear stain. Scale bars, 25 μm. (B) Quantification of the intensity of CD80-positive cells per field for each condition. Young; 8 - 9 weeks (n = 3) , Old; 30 months (n = 3). (C) Representative images of frozon sections of soleus muscle, stained with antibodies against CD206 for M2-type macrophages. DAPI was used as a nuclear stain. Scale bars, 25 μm. (D) Quantification of the intensity of CD206-positive cells per field for each condition. Young; 8 - 9 weeks (n = 3) , Aged; 30 months (n = 3). (B) and (D) were analyzed by One-way ANOVA and Tukey-Kramer test and represented as means ± SD. * $P < 0.05$, ** $P < 0.01$, and *** $P < 0.001$.

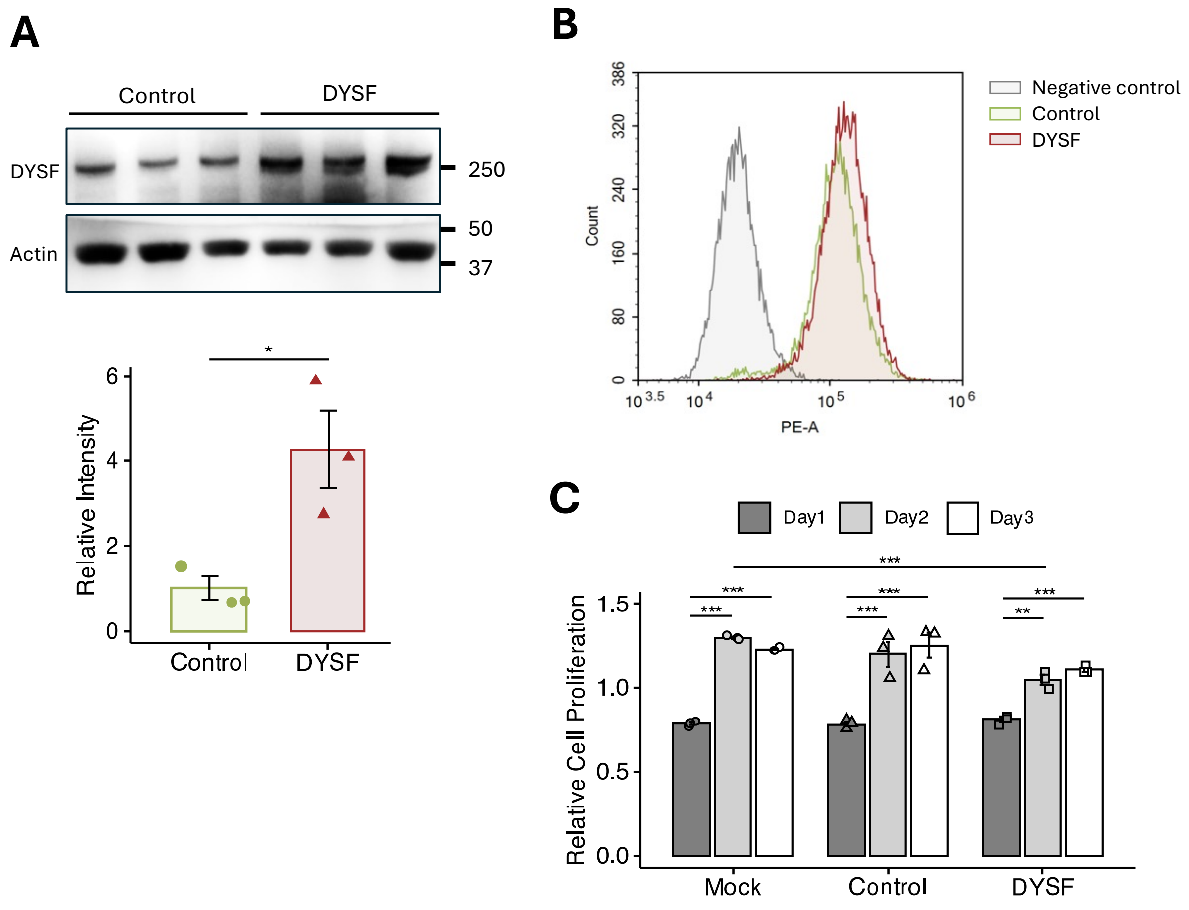


**Fig. S3 Cell Proliferation of DYSF Overexpression in Macrophages**

(A) Western blotting analysis of the expression of DYSF protein in J774A.1 cells transiently transfected with DYSF-3HA vector (DYSF) or pcDNA3.1 control vector (Control). β-actin (Actin) was used as a loading control (upper). The intensity of DYSF was normalized by Actin (lower). (B) Flow cytometry analysis of DYSF or Control vector-transfected J774A.1. Negative control, the negative control IgG antibody. (C) DYSF vector-transfected J774A.1 cells are effect cell proliferation compared to non-treated cells (Mock) or or Control vector-transfected cells (Control). (A) (lower) were analyzed by unpaired Student's t-test. (B) were analyzed by One-way ANOVA and Tukey-Kramer test. Figures [(A)(lower) and (C) ]represented as means ± SD. * $P < 0.05$, ** $P < 0.01$, and *** $P < 0.001$.

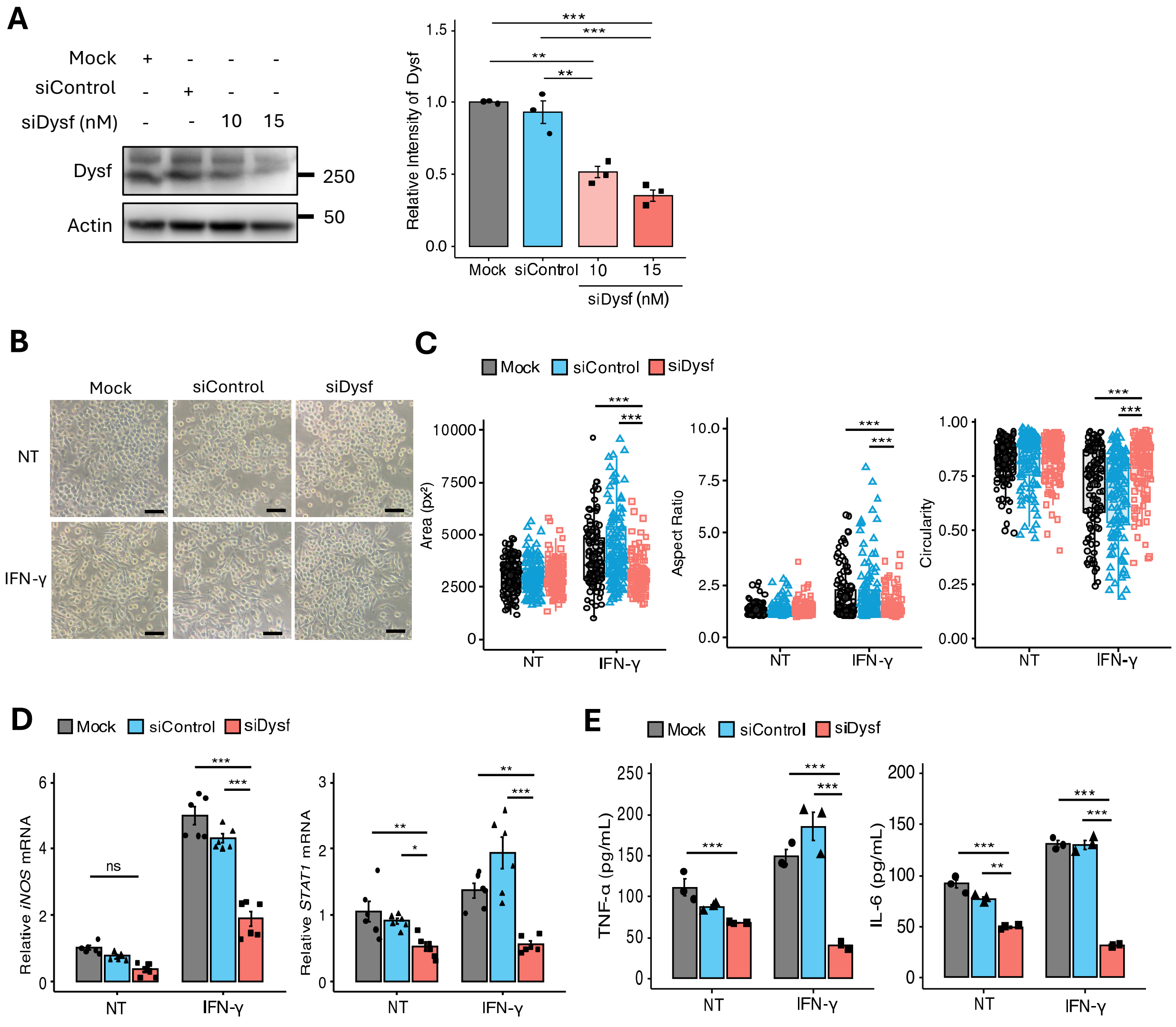


**Fig. S4 DYSF knockdown suppresses the charactaristics of IFN-γ-induced macrophages.**

(A) Expression of Dysf protein by western blotting in J774A.1 macrophages transiently transfected with *Dysf* siRNA (siDysf) or non-targeting siRNA (siControl), and non-transfected cells (Mock) (left). Density of the Dysf protein bands was normalized to β-actin (Actin) (right). n = 3. (B) Representative phase contrast images of J774A.1 transfected with siDysf, siControl and Mock were treated with or without 100 ng/mL IFN-γ for 24 hours. One hundred cells were counted for each condition. Scale bar: 50 µm. NT, Non-treated. (C) Quantification of pixel area, circularity and aspect ratio in siDysf and siControl -transfected J774A.1. One hundred cells were counted for each condition. (D) Expression of *iNOS* and *Stat1* mRNA in J774A.1 transfected with siDysf or siControl, and Mock was assessed by qPCR to confirm the characteristic of M1-type macrophages. n = 6. (E) The protein concentration of TNF-α and IL-6 in conditioned medium obtained from J774A.1 transfected with siDysf, siControl and Mock was assessed by ELISA. n = 3. (A) (right), (C), (D), and (E) were analyzed by One-way ANOVA and Tukey-Kramer test and represented as means ± SD. * $P < 0.05$, ** $P < 0.01$, and *** $P < 0.001$. ns, not significant.

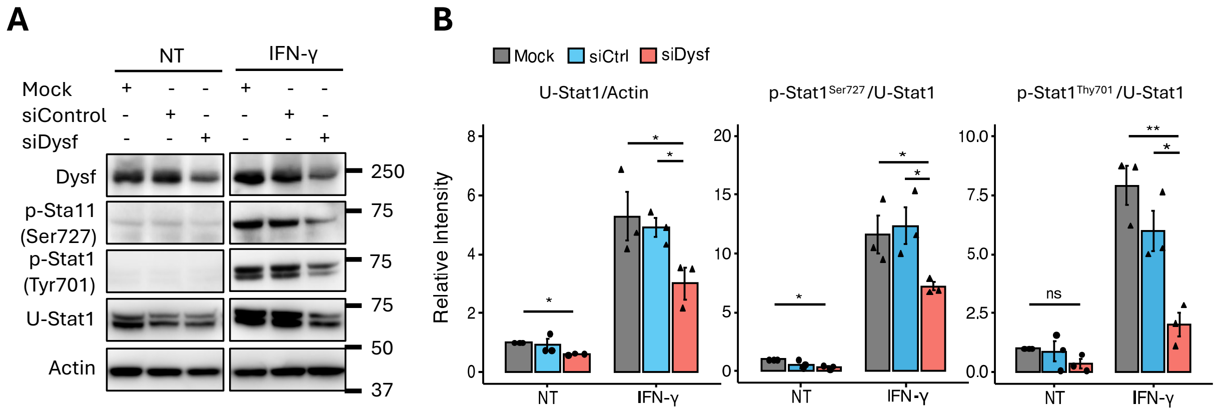


**Fig. S5 DYSF-knockdown Suppresses the Stat1 Activation in IFN-γ–induced Macrophages**

(A) The protein levels or the phosphorylation activation of Stat1 in J774A.1 transfected with siDYSF or siControl, and Mock were evaluated by western blotting. β-actin (Actin) was used as the loading control. NT, Non-treated. (B) Protein abundance of Stat1, Stat1 phospholated at Ser727 and Tyr701 in J774A.1 transfected with siDYSF or siControl, and Mock treated with or without IFN-γ (n = 3). Density of the protein bands was normalized to Actin or Stat1. (B) were analyzed by One-way ANOVA and Tukey-Kramer test and represented as means ± SD. * $P < 0.05$ and ** $P < 0.01$. ns, not significant.

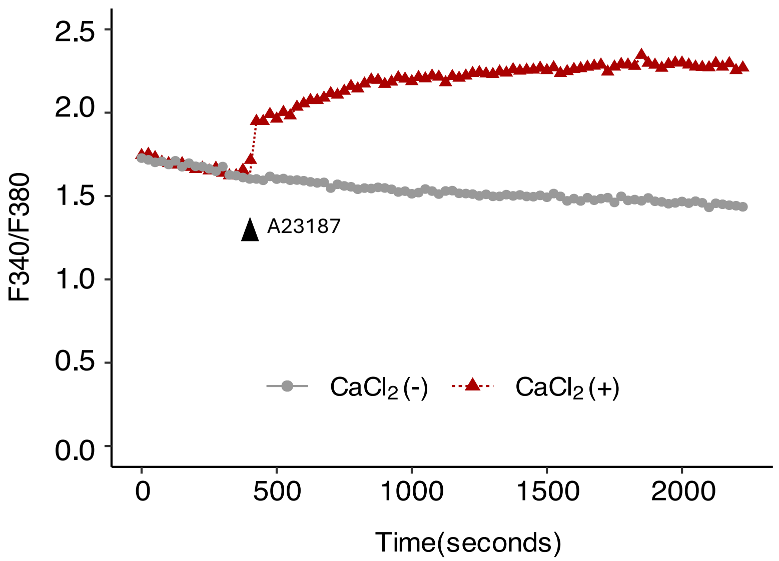


**Fig. S6 Effect of external calcium on calcium flux**

J774A.1 cells were loaded with Fura 2-AM (4 µg/ml) for 1 hour at 37°C in presence or absence of calcium. Calcium lonophore, A23187 (2 µM) was added during the measurement to detect the internal stores of calcium with Fura 2 ratio. We measured an excitation wavelength of 340/510 nm and an emission wavelength of 380/510 nm.

**Supplementary Tables**

**Table S1. List of antibodies in this study.**

| **Antibody** | **Reference** | **Use and Dilution** |
|---|---|---|
| STAT1 | Biolegend, 603701 | Western blotting (1:1000)<br>Immunofluorescence staining (1:100) |
| STAT1 Phospho (Ser727) | Biolegend, 686401 | Western blotting (1:1000) |
| STAT1 Phospho (Tyr701) | Cell Signaling Technology, 7649 | Western blotting (1:1000) |
| DYSF | Leica Biosystems, NCL-Hamlet | Western blotting (1:1000)<br>Immunofluorescence staining (1:100) |
| β-Actin | GeneTex, 14395-1-AP | Western blotting (1:5000) |
| GAPDH | Sigma-Aldrich, CB1001 | Western blotting (1:5000) |
| α-Tubulin | Biolegend, 627901 | Western blotting (1:1000) |
| Histon H3 (C-terminus) | Biolegend, 819411 | Western blotting (1:1000) |
| Myogenin | BD Biosciences, 556358 | Western blotting (1:1000)<br>Immunofluorescence staining (1:100) |
| MYH1 | Sigma-Aldrich, ZRB1214 | Immunofluorescence staining (1:100) |
| FITC-CD11b | Biolegend, 101205 | Immunofluorescence staining (1:100) |

**Table S2. List of primers used for quantitative RT-PCR (qPCR) in this study.**

| **Species** | **Gene** | **Forward (5' -> 3')** | **Reverse (5' -> 3')** |
|---|---|---|---|
| Mouse | GAPDH | TCCACCACCCTGTTGCTGTA | GACTTCAACAGCAACTCCCAC |
| Mouse | iNOS | ATCGACCCGTCCACAGTATG | GATGGACCCCAAGCAAGACT |
| Mouse | Arg-1 | GCACTGAGGAAAGCTGGTCT | GACCGTGGGTTCTTCACAAT |
| Mouse | TNF-α | GAACTGGCAGAAGAGGCACT | AGGGTCTGGGCCATAGAACT |
| Mouse | STAT1 | GTTCCGACACCTGCAACTGAA | AGAGGTGGTCTGAAAGGGAAC |
| Human | GAPDH | CATGTTCCAATATGATTCCACC | CTCCACGACGTACTCAGCG |
| Human | DYSF | AAGGATGCCTTCTGGAGG | GGTCATTTCCAGCTTGCC |
